# Machine learning-based prediction of memory requirements for metagenomic assembly in high-performance computing environments

**DOI:** 10.64898/2026.05.12.724571

**Authors:** Santiago Sanchez, Santino Faack, Martin Beracochea, Robert D. Finn, Björn Grüning, Bérénice Batut, Paul Zierep

## Abstract

Metagenomic assembly can be a computationally intensive step in microbiome analysis, with memory requirements that vary widely depending on input data characteristics. In workflow systems like Galaxy and large-scale platforms like MGnify, which run thousands of heterogeneous jobs, inaccurate memory allocation drives job failures and costly retries when underestimated, and reduces throughput when overestimated. Current approaches rely primarily on heuristic rules based on input file size or sample metadata, which often fail to generalize across diverse datasets.

In this study, we present a machine learning-based framework for predicting memory requirements of metagenomic assembly using metaSPAdes. We analyzed 300 assembly jobs from diverse biomes and evaluated 18 predictive models using combinations of input file size, biome classification, and sequence-derived k-mer features. K-mer profiles were computed from raw sequencing data and summarized into statistical descriptors capturing sequence complexity and diversity. Model performance was assessed using both conventional regression metrics and a production-oriented cost function that accounts for retry policies and resource waste in high-performance computing environments.

Our results show that machine learning models can outperform commonly used heuristics. In particular, models incorporating biome information achieved the best performance and can be tuned to favor conservative predictions that reduce job failure rates. Simpler models based solely on input file size also performed competitively, offering a practical alternative for systems with limited feature availability. When evaluated under realistic workload distributions, predictive approaches reduced total memory waste by several million gigabyte-hours per 1,000 jobs compared to static allocation strategies.

These findings demonstrate that data-driven resource prediction can substantially improve efficiency in metagenomic workflows. The proposed framework is adaptable to different computational environments and provides a foundation for integrating predictive resource allocation into large-scale bioinformatics platforms beyond Galaxy.

## Background

Advances in high-throughput sequencing have revolutionized microbiome research, allowing scientists to study microbial communities without the need for cultivation. Among these advances, whole metagenome shotgun sequencing (WMGS) has gained widespread use, as it directly sequences all the genomes in a sample and offers simultaneous insights into both taxonomic diversity and functional gene content (Quince et al. 2017). Metagenomic assembly is a key step in processing generated data, e.g. for reconstructing genomes. However, it remains computationally demanding, particularly in terms of memory consumption (Meyer et al. 2022; Sun et al. 2022). Assemblers such as metaSPAdes (Nurk et al. 2017) rely on de Bruijn graph–based approaches, in which memory usage is largely determined by the number of distinct k-mers stored as graph nodes, a quantity that scales according to sequencing depth and community complexity (Rødland 2013; Chikhi et al. 2016; Salikhov et al. 2014). As a result, predicting resource requirements for metagenomic assembly remains a significant challenge.

In large-scale bioinformatics platforms and high-performance computing (HPC) environments, inefficient resource allocation can have substantial consequences. Underestimation of memory requirements leads to job failures and repeated executions, increasing overall computational cost and delaying analyses. Conversely, overestimation results in underutilized resources, reducing system throughput and efficiency. These challenges are particularly relevant in workflow management systems such as Galaxy (The Galaxy Community 2024) and large-scale infrastructures like MGnify (Richardson et al. 2023), where thousands of jobs are executed routinely across heterogeneous datasets. Improving resource allocation efficiency not only enhances HPC performance but also reduces environmental impact by lowering unnecessary energy consumption in compute infrastructure.

Current approaches to resource allocation are typically heuristic-based. For example, memory assignment may be determined using simple rules derived from input file size or predefined mappings based on sample metadata such as biome classification. While these heuristics are straightforward to implement, they often fail to capture the complexity of metagenomic data. A recent study suggests that sequence-derived features, such as k-mer profiles, provide a more informative representation of dataset complexity and may improve the prediction of computational requirements (Belmann et al. 2025). Machine learning approaches offer a promising framework for integrating such features with metadata and input size to generate more accurate and adaptive resource estimates.

In this study, 300 metagenomic assembly jobs across diverse biomes were used to evaluate 18 predictive models and identify the best approach for estimating memory requirements of metaSPAdes (v3.15.3) based on input data size, biome classification, and/or k-mer profile statistics. Although there are multiple tools available to count kmers frequencies like jellyfish (Marçais and Kingsford 2011a) and KMC (Kokot et al. 2017) we are not aware of tools able to compute summary statistics like alpha diversity directly from kmer profiles. To address this gap, we developed *kmer2stats (Faack and Zierep 2026)*, a dedicated tool for deriving statistical features from k-mer abundance profiles. This tool was integrated into a FAIR Galaxy workflow that enables direct processing from SRA accessions, covering all steps from data retrieval and read processing to k-mer counting and feature extraction.

The performance of these predictive models on the 300 metagenomic assembly jobs was compared against default heuristic strategies used in MGnify and Galaxy. Model performance was assessed using conventional metrics, including root mean squared error (RMSE) and coefficient of determination (R^2^). However, high-performance computing (HPC) environments such as MGnify and Galaxy implement specific job execution policies, including retry mechanisms for jobs that fail due to insufficient memory allocation. To account for this behavior, we introduce a novel scoring function that estimates total memory consumption over time under retry-based execution. This metric provides a more realistic representation of resource usage than standard evaluation measures. Comparison with the novel scoring function indicates that traditional metrics may overestimate predictive performance under realistic, retry-based execution scenarios.

## Materials and methods

### Training and validation dataset

We compiled a dataset of 9,102 metagenomics assembly jobs (metaSPAdes v3.15.3) processed by EMBL-EBI using the MGnify metagenome assembly pipeline (https://github.com/ebi-metagenomics/miassembler). The analysed sequence datasets were sourced from the European Nucleotide Archive (ENA), and the resulting assemblies were deposited back into ENA following processing. Pipeline execution metadata were recorded during processing on the EMBL-EBI High Performance Computing (HPC) infrastructure, which provides access to high-memory compute nodes.

The dataset included fields for primary accession, sample biome, peak memory usage (GB), assembler, and assembler version. From these, 300 jobs were randomly selected as a subsample for model development, comprising 30 samples from each major biome (Supplementary Table 1) to ensure balanced representation across biomes.

To characterize the k-mer profile of the selected jobs, raw sequencing reads were retrieved through the INSDC archives using accession metadata obtained via the EMBL-EBI API (Madeira et al. 2019). Raw reads were downloaded, and k-mers were computed using Jellyfish (v2.3.0) (Marçais and Kingsford 2011b) with a k-mer length of 10 and otherwise default parameters. From these k-mer counts, 39 statistical features (Table 2), including count-based and diversity metrics, were derived using kmer2stats (v1.0.1) (Faack 2026) (Supplementary Table 2). A Galaxy workflow that reproduces the full analysis, including all processing steps and computation of k-mer statistics from sample SRA accessions, is available at: https://usegalaxy.eu/u/paulzierep/w/srr-to-kmer-statistics.

**Table 1.**
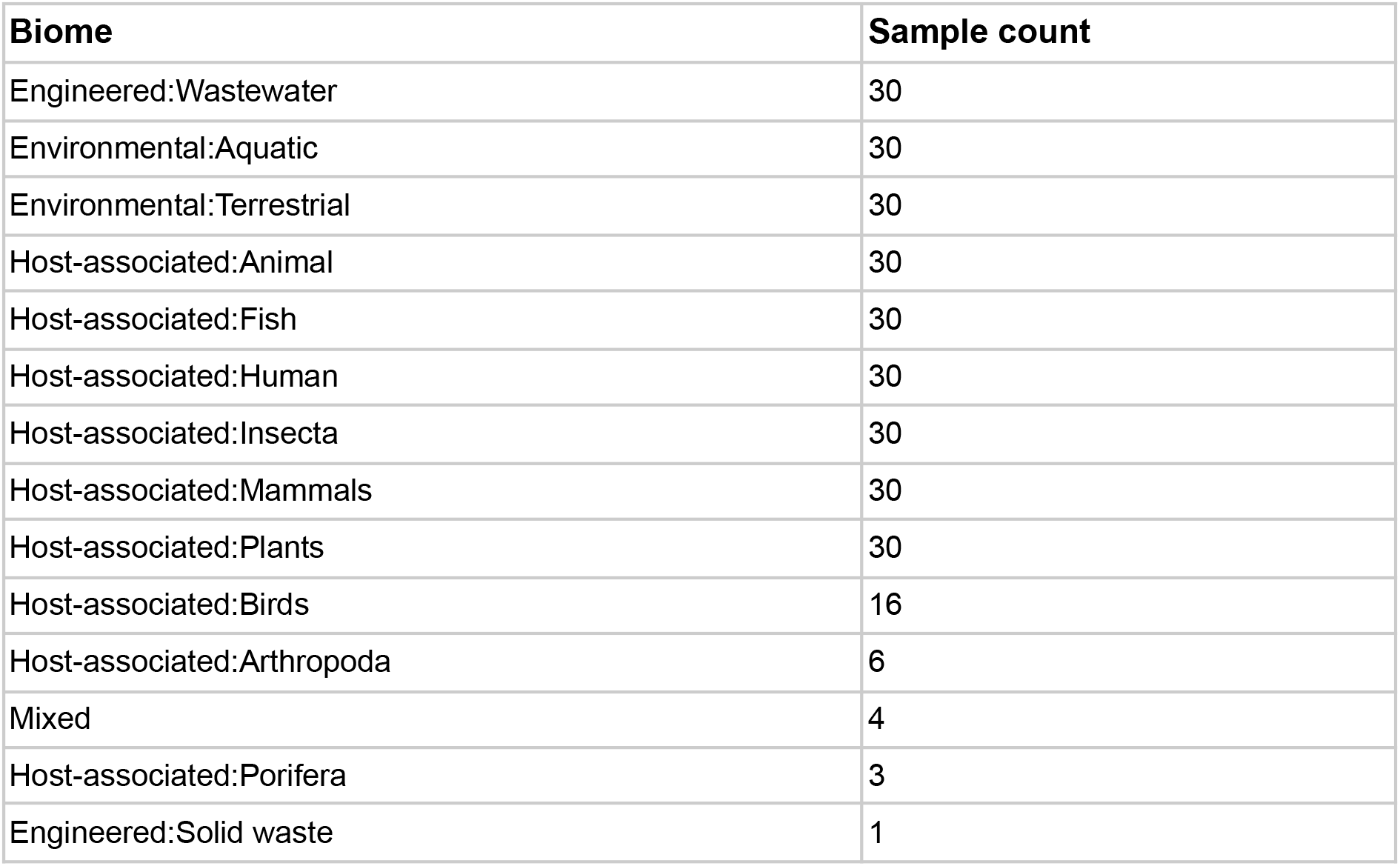
Distribution of sample biomes in this study based on GOLD classification.

**Table 2.** Overview of k-mer-derived statistical features generated by kmer2stats. The tool transforms raw k-mer counts (e.g., from Jellyfish) into 39 summary statistics describing abundance, diversity, sparsity, and distributional properties of k-mer profiles.

| Feature group | Example features | Description |
| --- | --- | --- |
| Count statistics | total k-mer counts<br>unique k-mers<br>non-zero k-mers | Basic abundance and complexity measures derived from k-mer frequency tables |
| Frequency distribution metrics | mean<br>median<br>variance<br>standard deviation | Describes the distribution of k-mer counts across all observed k-mers |
| Sparsity measures | zero proportion<br>richness | Quantifies proportion of absent vs. observed k-mers and overall sequence sparsity |
| Diversity indices | Shannon entropy<br>Simpson diversity | Captures k-mer diversity and evenness in the sample |
| Dominance metrics | max frequency | Measures whether a few |
|  | top-k-mer proportion | number of k-mers dominate the profile |
| Complexity indicators | normalized entropy<br>complexity score | Reflects sequence complexity based on k-mer distribution shape |
| Skewness-related features | skewness<br>kurtosis | Captures asymmetry and tail heaviness of k-mer frequency distribution |
| Coverage-related features | estimated coverage proxy<br>abundance concentration | Approximates unevenness in sequencing depth across k-mers |

**Table 2.** Comparison of predictive models and heuristic approaches for memory allocation in metaSPAdes workflows (sorted by R^2^ values). Models are evaluated in terms of predictive accuracy (R^2^, RMSE) and downstream computational cost under a retry-aware HPC execution policy. Waste is reported as mean ± uncertainty per 1,000 jobs in million GB·h.

| Model | Features | R<sup>2</sup> | RMSE | Waste per 1,000 jobs (million GB·h) |
| --- | --- | --- | --- | --- |
| Random Forest | K-mers + File size + Biome | **0.68** | 31.05 | 3.2 ± 1.51 |
| Random Forest | K-mers + File size | 0.65 | 32.45 | 3.17 ± 1.56 |
| Random Forest | Biome | 0.47 | 39.98 | **2.45 ± 0.79** |
| LinearRegression | Biome | 0.47 | 40.03 | 2.47 ± 0.8 |
| Gaussian Processes (Quantile 0.9) | Biome | 0.46 | 40.23 | 2.64 ± 0.96 |
| Gaussian Processes (Quantile 0.99) | Biome | 0.45 | 40.72 | 2.88 ± 1.15 |
| LinearRegression | K-mers + File size + Biome | 0.29 | 46.27 | 4.17 ± 2.18 |
| Gaussian Processes (Quantile 0.9) | K-mers + File size + Biome | 0.21 | 48.59 | 2.49 ± 0.9 |
| LinearRegression | File size | 0.18 | 49.63 | 3.53 ± 1.32 |
| Gaussian Processes (Quantile 0.9) | K-mers + File size | 0.11 | 51.57 | 2.69 ± 0.99 |
| Random Forest | File size | 0.02 | 54.10 | 4.51 ± 2.06 |
| LinearRegression | K-mers + File size | -0.03 | 55.55 | 4.42 ± 2.11 |
| Gaussian Processes (Quantile 0.99) | K-mers + File size + Biome | -0.53 | 67.82 | 3.35 ± 1.26 |
| Gaussian Processes (Quantile 0.9) | File size | -0.70 | 71.38 | 20.69 ± 17.91 |
| Gaussian Processes (Quantile 0.99) | K-mers + File size | -0.77 | 72.89 | 2.88 ± 0.71 |
| Gaussian Processes (Quantile 0.99) | File size | -1.01 | 77.65 | 5.23 ± 1.67 |
| MGNify Heuristic default | Heuristic | -1.46 | 85.84 | 3.27 ± 1.01 |
| Galaxy Heuristic default | Heuristic | -18.19 | 239.88 | 9.03 ± 1.51 |

### Machine learning models and baseline comparisons

To predict the memory consumption of metaSPAdes using the 39 different k-mer statistics, input file size, and/or sample biome classification, three machine learning models were implemented using scikit-learn (v1.8.0) (Pedregosa et al. 2011): (i) random forest regressor,(ii) linear regression, (iii) a custom Gaussian process regressor (based on scikit-learn) incorporating preprocessing steps within a modelling pipeline, optional log-transformation of the target variable to address skewed memory usage distribution, predictive uncertainty estimates, and quantile-based (0.9 and 0.99) predictions, enabling memory allocation based on upper bound estimates to minimize failures due to underestimation. For each model, feature sets were combined to assess their contribution to predictive performance.

In addition to the machine learning models, two heuristics currently used in production systems were also evaluated: (i) Galaxy job memory heuristic, a rule-based approach assigning memory in steps based on the input file size, (ii) MGnify memory estimation heuristic, a biome-specific mapping-table (Supplementary Table 3). These heuristics were compared against the machine learning models to assess their relative accuracy in predicting memory requirements. In total, 18 models were evaluated on out-of-fold predictions generated by five-fold shuffled cross-validation on the 300 assembly job data.

### Model evaluation

Model accuracy was measured using root mean squared error (RMSE) and coefficient of determination (R^2^). While these metrics quantify the overall prediction accuracy, they do not capture the operational impact of allocation errors. In practice, these two errors are asymmetric in cost. Under-prediction wastes the resources of every failed attempt before a larger retry succeeds, whereas over-prediction wastes only the unused portion of a single allocation. To account for these effects, a cost function was defined that estimates total resource waste under a retry policy. If a job fails due to insufficient memory, it is retried with incrementally larger allocations until successful. The cost function quantifies the total resource consumption, including both failed attempts and unused memory in successful runs.

Under the retry policy

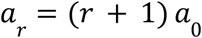

where *a*_*0*_ is the initial allocation and *r* is the retry index, the job repeats until the allocated memory exceeds the true peak usage (*y*). The number of failed attempts is

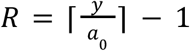

The total resource cost per job is

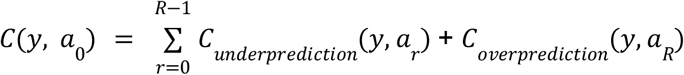

Failed attempts consume compute resources before termination. If attempt *r* allocates *a*_*r*_ *< y*, the job runs for a fraction *a*_*r*_*/y* of its true runtime *t*:

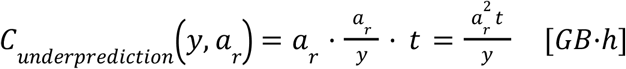

On the successful attempt, unused reserved memory represents wasted capacity for the full runtime:

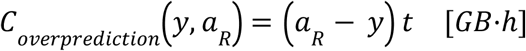

Performances of each model were evaluated using Monte Carlo simulations designed to reflect the memory usage distribution observed in the production workload. The production workload distribution was estimated from the assembly jobs dataset, where peak memory usage ranged from <1 GB to several hundred GB and exhibited a strongly right-skewed distribution. Most jobs required modest memory allocations, while a smaller number of large assemblies consumed substantially more.

Since the evaluation dataset used for cross-validation does not necessarily match the production workflow distribution, each evaluation job was assigned a weight reflecting its frequency in production:

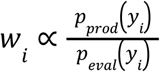

where p_prod_(y) and p_eval_(y) are the production workload density and the evaluation dataset density, respectively.

Densities were estimated using histograms with 60 log-spaced bins across the combined memory range (∼1–300 GB) to preserve resolution across the wide range of memory values. Jobs receive higher weights when their memory usage is common in production but rare in the evaluation set, ensuring the simulation reflects real production frequencies.

For each model, 4,000 Monte Carlo iterations were performed. In each iteration 1,000 jobs were sampled with replacement from the evaluation dataset, with sampling probabilities proportional to the production-distribution weights (*w*_*i*_). For each sampled job, the cost function was evaluated using the model’s predicted allocation (*a*_0_). When production wall-clock times were available, they were used directly. Otherwise, the runtime of a failed attempt was approximated as (*a*_*r*_ / *y*).

The simulations produced a distribution of total memory waste across iterations for each model. Results were summarised as the posterior mean and 95% interval (2.5–97.5 percentiles) and reported as GB·h of wasted resources per 1,000 jobs. This production-weighted Monte Carlo estimate served as the primary metric for comparing model performance.

## Results

The resource requirements of metaSPAdes (v3.15.3) were estimated using both HPC heuristics and predictive models trained on different feature sets, by assessing the agreement between predicted peak memory and measured peak memory (Figure 1 and Table 2). The MGnify heuristic showed substantially better agreement with the ideal prediction than the Galaxy heuristic (R^2^ = -1.46 and R^2^ = -18.19, respectively).

**Figure 1.**
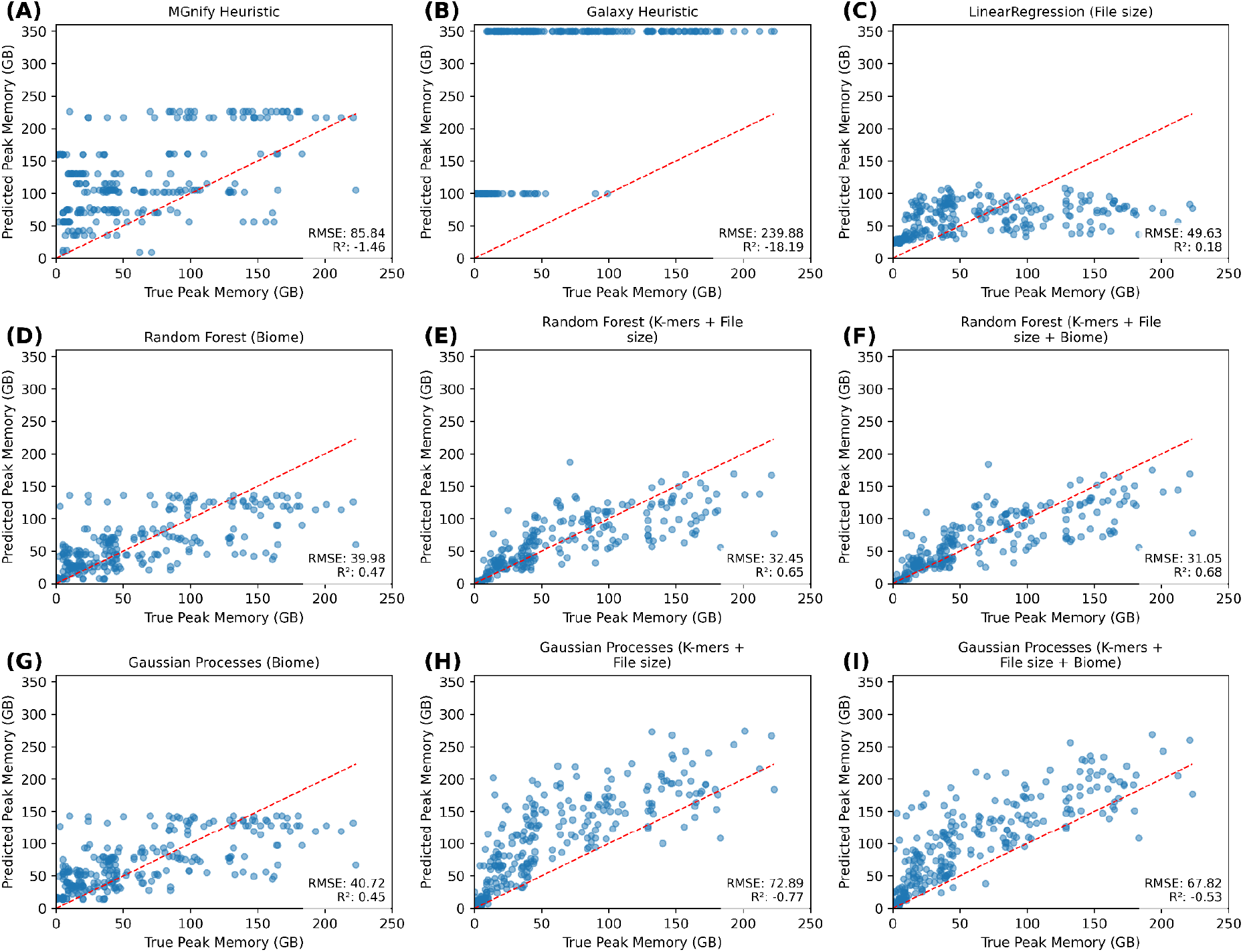
Resource prediction using different heuristics and models trained on different feature sets. For clarity only 9 models are shown. Metrics for all 18 models are listed in Table 3. Predicted (y-axis) vs. true (x-axis) peak memory (GB) for 300 metaSPAdes assembly jobs spanning multiple biomes, evaluated under 5-fold cross-validation. Each blue point is one job; the dashed red diagonal marks perfect prediction. Root-mean-squared error (RMSE) and coefficient of determination (R^2^) for each predictor are inset. **(A)** MGnify heuristic using the biome mapping table. **(B)** Galaxy heuristic using the rule-based approach. **(C)** Linear model using only input file size. **(D)** Random Forest trained on k-mer statistics and input file size. **(E)** Gaussian Process trained on k-mer statistics and input file size. **(F)** Gaussian Process trained on k-mer statistics, input file size, and biome.

Overall, machine learning models trained on input features such as k-mer statistics, file size, and biome information outperformed both heuristic approaches. The best-performing model was a Random Forest trained on the full feature set (k-mers + file size + biome), achieving an R^2^ of 0.68. Models using only biome information also showed reasonable predictive performance, with both Random Forest and Linear Regression achieving R^2^ ≈ 0.47.

Since biome annotation requires manual input, it is relevant to evaluate models based solely on automatically derivable features. In this context, the Random Forest model trained on k-mers and file size alone still achieved strong performance (R^2^ = 0.65). The simplest model, Linear Regression using only file size as input, does not require k-mer computation or manual annotation and achieved moderate predictive performance (R^2^ = 0.18).

Both Galaxy and MGnify implement a retry policy within their workflow execution settings. This policy follows a simple but impactful rule: if a job fails due to insufficient memory, the allocated memory is doubled for the subsequent attempt. As a consequence, underestimation of memory requirements propagates into increased resource consumption over repeated executions (Figure 2A). To capture this effect, predictive performance was reassessed under the retry policy, reporting memory costs in million GB·h per 1,000 jobs (Figures 2B).

**Figure 2.**
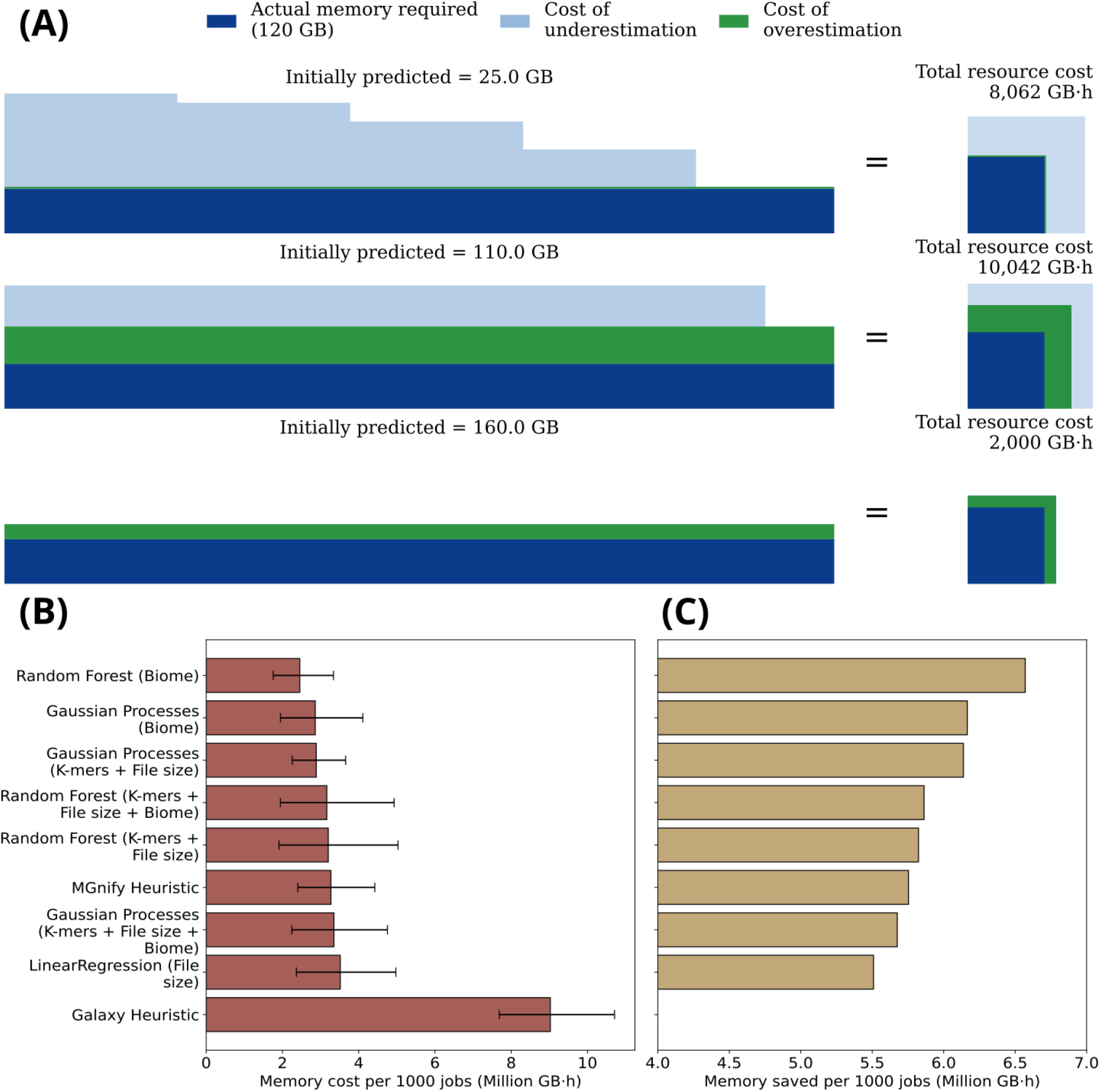
Cost-aware memory allocation for metagenome assembly. **(A)** Representation of how memory–time waste accumulates for a hypothetical assembly job whose true peak memory is 120 GB and whose true wall time is 50 h, evaluated under three illustrative initial allocations (a0), using a standard memory allocation retry policy (scales linearly on successive attempts: a0, 2·a0, 3·a0, …). Each attempt is drawn as a rectangle whose width is wall time (h) and height is memory allocation (GB), so its area is memory–time consumption (GB·h). Light-blue rectangles stacked above the success rectangle are failed attempts; Successful attempts are subdivided: (1) the solid dark-blue rectangle at the bottom of every row marks the needed resources (120 GB × 50 h = 6,000 GB·h) the job actually consumes on success and (2) the green band shows the wasted memory–time allocation at successful job attempts (allocated resources minus needed resources). The blue square at the right of each row reduces the cumulative memory–time cost to an equivalent area, drawn on a common scale across the three rows for visual comparison. *Top*: a severe under-estimate (a0 = 25 GB) cascades through four failures (25, 50, 75, 100 GB) before succeeding at 125 GB and accumulates 8,062 GB·h of waste. *Middle*: a near-miss (a0 = 110 GB) fails once close to the end of its wall time and then over-allocates at 220 GB on retry, costing 10,042 GB·h, the worst of the three despite missing the truth by only 10 GB. *Bottom*: a modest over-estimate (a0 = 160 GB) succeeds on the first attempt for 2,000 GB·h, five times less than the near-miss, even though its absolute deviation from the truth (40 GB) is larger. **(B)** Mean predicted memory cost per 1,000 jobs (million GB·h) for each predictor; error bars are 95% CI from 4,000 weighted Monte Carlo iterations. **(C)** Memory saved per 1,000 jobs relative to the baseline predictor (Galaxy heuristic, 9.03 million GB·h). Predictors are ordered by mean cost.

Under these conditions, the best-performing predictor was the Random Forest model trained only on biome information, with a cost of 2.456 ± 0.796 Million GB·h per 1,000 jobs. A comparable performance was observed for the Gaussian model trained on biome features (2.639 ± 0.966 million GB·h) as well as for the Random Forest models using k-mer and file-size features (3.170 ± 1.560 million GB·h) or all feature sets combined (3.201 ± 1.508 million GB·h). The best model using only automatically derivable features was the Gaussian Processes model trained on k-mer statistics and file size, evaluated under both quantiles. The quantile 0.9 configuration achieved 2.69 ± 0.99 million GB·h per 1,000 jobs, while the quantile 0.99 configuration resulted in 2.88 ± 0.71 million GB·h per 1,000 jobs.

Several additional models achieved intermediate performance. For example, the MGnify heuristic resulted in 3.272 ± 1.013 million GB·h, while linear regression models ranged from 2.465 ± 0.800 million GB·h (biome only) to 4.416 ± 2.110 Million GB·h (k-mers and file size). Models based solely on file size also performed moderately, with 3.531 ± 1.326 million GB·h (linear regression).

This metric demonstrates that models such as Gaussian Processes, as well as the MGnify heuristic, which may appear less competitive under traditional predictive metrics, can achieve strong performance when evaluated under realistic HPC retry logic.

Consequently, substantial resource savings can be achieved by adopting predictive or even simple linear allocation strategies (Figure 2C). In high-throughput settings, the Random Forest model trained on biome information reduces memory usage by more than 6.58 million GB·h per 1,000 jobs compared to the Galaxy heuristic (9.04 ± 1.52 million GB·h). The best-performing model using only automatically derivable features (Gaussian Processes trained on k-mer statistics and file size) also achieves substantial savings, reducing memory usage by more than 6 million GB·h per 1,000 jobs relative to the Galaxy heuristic.

Similarly, the MGnify heuristic provides a reduction of approximately 5.8 million GB·h per 1,000 jobs, while even simple linear models trained solely on file size achieve savings exceeding 5.5 million GB·h per 1,000 jobs. Overall, these results highlight that both predictive and heuristic approaches can significantly improve resource efficiency under retry-aware execution policies, with performance differences becoming most apparent at scale.

## Discussion and Outlook

Efficient allocation of computational resources is a critical challenge in large-scale metagenome assembly workflows. Our results demonstrate that both predictive models and existing HPC heuristics differ substantially in their ability to approximate true memory requirements and, more importantly, in their downstream cost under realistic execution policies.

Under standard evaluation based on peak-memory prediction accuracy (Figure 1 and Table 2), machine learning models consistently outperformed rule-based heuristics. The MGnify heuristic showed improved agreement with observed memory usage compared to the Galaxy heuristic (R^2^ = −1.46 vs. −18.19), although both methods performed poorly in absolute terms. In contrast, all learned models achieved substantially higher predictive accuracy, with the best-performing Random Forest model (k-mers + file size + biome) reaching R^2^ = 0.68 and the lowest RMSE (31.05 GB).

Importantly, even restricted feature sets provided strong predictive power. Models trained solely on biome information achieved moderate performance (R^2^ ≈ 0.47), indicating that environmental context alone is partially informative of computational requirements. When only automatically derivable features were considered, the Random Forest model using k-mer statistics and file size still maintained strong accuracy (R^2^ = 0.65), approaching that of the full feature set. In contrast, the simplest model based only on file size yielded substantially lower performance (R^2^ = 0.18), highlighting the limitations of purely size-based estimation.

When incorporating HPC retry logic, the relative performance landscape changes substantially. Under a memory-doubling retry policy, predictive accuracy translates directly into differences in cumulative computational cost (Figure 2). Here, the Galaxy heuristic performed worst overall (9.03 ± 1.51 million GB·h per 1,000 jobs). The MGnify heuristic reduced this cost considerably (3.27 ± 1.01 million GB·h), but still remained above most predictive models.

Among learned approaches, the best-performing model under retry-aware evaluation was the Random Forest trained on biome features (2.45 ± 0.79 million GB·h), closely followed by Gaussian Process models using biome or automatically derived features (≈2.6–2.9 million GB·h depending on quantile).

Crucially, models relying only on automatically derivable features also performed strongly under this cost-based metric. The Gaussian Process model using k-mer statistics and file size achieved 2.69 ± 0.99 million GB·h (0.9 quantile) and 2.88 ± 0.71 million GB·h (0.99 quantile), placing it among the most efficient non-biome-dependent approaches. Even simple linear models and MGnify-style heuristics achieved competitive performance in the 2.5-4.5 million GB·h range, demonstrating that useful efficiency gains are achievable without complex feature engineering.

These results highlight an important distinction between predictive accuracy and operational efficiency. While R^2^ and RMSE capture agreement to the observed peak memory, they do not directly reflect system-level cost under retry-aware execution. In this context, even models with moderate predictive accuracy can yield substantial reductions in total resource consumption compared to static heuristics.

Overall, our findings show that significant efficiency gains, exceeding 6 million GB·h per 1,000 jobs relative to the Galaxy heuristic, can be achieved through learned allocation strategies. Importantly, competitive performance is possible even when restricting models to automatically derivable features, making these approaches practical for large-scale deployment.

However, the retry logic introduced here warrants further investigation to enable more accurate estimation of memory costs under realistic HPC execution behaviour. In the current formulation, we assume a linear relation between allocated memory and job failure time (Figure 2A), but we do not model or observe the exact point at which a job fails during execution. In practice, job failure occurs at an unknown and potentially variable stage of execution, which may depend on multiple factors. Consequently, the true failure time is not directly observable, and this uncertainty may significantly influence the resulting memory–time cost estimates. A further limitation is that our cost estimates exclude the overhead of computing k-mer statistics, which should be considered when evaluating the net efficiency gains. In addition, the retry-aware framework evaluated here does not incorporate the internal retry and checkpointing implemented in metaSPAdes. In practice, metaSPAdes can recover from certain failed stages and resume execution without restarting the entire process. However, such behaviour is not currently supported by Galaxy or Nextflow, which typically treat failed jobs as task failures that require full resubmission. As a result, the cost model presented here does not consider assembler-native recovery strategies.

Consequently, we propose to consider standard evaluation based on peak-memory prediction accuracy as well as HPC retry logic to decide for ideal models for memory assignment.

From an implementation perspective, these findings have direct implications for workflow systems such as MGnify and Galaxy. MGnify is well-positioned to integrate predictive models into its scheduling system. The assembly pipeline is implemented in Nextflow, which allows workflow resource requirements to be dynamically assigned. This makes metadata-based predictive models practical candidates for large-scale resource allocation.

In contrast, current limitations within Galaxy’s Total Perspective Vortex (TPV) (https://github.com/galaxyproject/total-perspective-vortex), the system that assigns resources for Galaxy jobs prevent the direct execution of machine learning models during scheduling, due to limited support for runtime evaluation of predictive models and security considerations associated with user-provided inputs. Consequently, we implemented a linear model based on file size within Galaxy’s TPV, as this approach can be expressed as a simple formula compatible with the system’s configuration.

Overall, our results demonstrate that substantial resource savings, on the order of millions of gigabyte-hours per 1,000 jobs, can be achieved through predictive or semi-predictive allocation strategies. Future work should focus on improving the portability and integration of such models across workflow platforms.

## Declarations

### Data and Code Availability

All code and analysis scripts, along with processed data required to reproduce the results, are available in the following GitHub repository: https://github.com/usegalaxy-eu/metaspades-memory-prediction

kmer2stats (https://github.com/SantaMcCloud/kmer2stats) is available via PyPI (https://pypi.org/project/kmer2stats) and Bioconda (https://anaconda.org/bioconda/kmer2stats), and is also provided as a Galaxy tool wrapper (toolshed.g2.bx.psu.edu/repos/iuc/kmer2stats/kmer2stats/1.0.1+galaxy1 available on usegalaxy.eu).

The Galaxy workflow used for k-mer generation from SRA accessions is available at: https://usegalaxy.eu/u/paulzierep/w/srr-to-kmer-statistics

The MGnify assembly pipeline is available in the following GitHub repository: https://github.com/EBI-Metagenomics/miassembler and Workflowhub link: https://workflowhub.eu/workflows/2171

### Funding

German Federal Ministry of Education and Research, BMBF [031 A538A de.NBI-RBC]. Ministry of Science, Research and the Arts Baden-Württemberg (MWK) within the framework of LIBIS/de.NBI Freiburg.

de.NBI Cloud within the German Network for Bioinformatics Infrastructure (de.NBI) and ELIXIR-DE (Forschungszentrum Jülich and W-de.NBI-001, W-de.NBI-004, W-de.NBI-008, W-de.NBI-010, W-de.NBI-013, W-de.NBI-014, W-de.NBI-016, W-de.NBI-022).

The French Institute of Bioinformatics (IFB) was founded by the Future Investment Program subsidized by the National Research Agency, number ANR-11-INBS-0013.

European Molecular Biology Laboratory core funds.

## References

Belmann, Peter, Benedikt Osterholz, Nils Kleinbölting, Alfred Pühler, Andreas Schlüter, and Alexander Sczyrba. 2025. “Metagenomics-Toolkit: The Flexible and Efficient Cloud-Based Metagenomics Workflow Featuring Machine Learning-Enabled Resource Allocation.” NAR Genomics and Bioinformatics 7 (3): lqaf093. 10.1093/nargab/lqaf093.

Chikhi, Rayan, Antoine Limasset, and Paul Medvedev. 2016. “Compacting de Bruijn Graphs from Sequencing Data Quickly and in Low Memory.” Bioinformatics 32 (12): i201–8. 10.1093/bioinformatics/btw279.

Faack, Santino. 2026. Kmer2stats. Zenodo, released April 27. 10.5281/zenodo.19828576.

Faack, Santino, and Paul Zierep. 2026. Kmer2stats. Zenodo, released May 4. 10.5281/zenodo.20026288.

Kokot, Marek, Maciej Długosz, and Sebastian Deorowicz. 2017. “KMC 3: Counting and Manipulating k-Mer Statistics.” Bioinformatics 33 (17): 2759–61. 10.1093/bioinformatics/btx304.

Madeira, Fábio, Young Mi Park, Joon Lee, et al. 2019. “The EMBL-EBI Search and Sequence Analysis Tools APIs in 2019.” Nucleic Acids Research 47 (W1): W636–41. 10.1093/nar/gkz268.

Marçais, Guillaume, and Carl Kingsford. 2011a. “A Fast, Lock-Free Approach for Efficient Parallel Counting of Occurrences of k-Mers.” Bioinformatics 27 (6): 764–70. 10.1093/bioinformatics/btr011.

Marçais, Guillaume, and Carl Kingsford. 2011b. “A Fast, Lock-Free Approach for Efficient Parallel Counting of Occurrences of k-Mers.” Bioinformatics 27 (6): 764–70. 10.1093/bioinformatics/btr011.

Meyer, Fernando, Adrian Fritz, Zhi-Luo Deng, et al. 2022. “Critical Assessment of Metagenome Interpretation: The Second Round of Challenges.” Nature Methods 19 (4): 429–40. 10.1038/s41592-022-01431-4.

Nurk, Sergey, Dmitry Meleshko, Anton Korobeynikov, and Pavel A. Pevzner. 2017. “metaSPAdes: A New Versatile Metagenomic Assembler.” Genome Research 27 (5): 824–34. 10.1101/gr.213959.116.

Pedregosa, Fabian, Gaël Varoquaux, Alexandre Gramfort, et al. 2011. “Scikit-Learn: Machine Learning in Python.” Journal of Machine Learning Research 12 (85): 2825–30.

Quince, Christopher, Alan W. Walker, Jared T. Simpson, Nicholas J. Loman, and Nicola Segata. 2017. “Shotgun Metagenomics, from Sampling to Analysis.” Nature Biotechnology 35 (9): 833–44. 10.1038/nbt.3935.

Richardson, Lorna, Ben Allen, Germana Baldi, et al. 2023. “MGnify: The Microbiome Sequence Data Analysis Resource in 2023.” Nucleic Acids Research 51 (D1): D753–59. 10.1093/nar/gkac1080.

Rødland, Einar Andreas. 2013. “Compact Representation of K-Mer de Bruijn Graphs for Genome Read Assembly.” BMC Bioinformatics 14 (1): 313. 10.1186/1471-2105-14-313.

Salikhov, Kamil, Gustavo Sacomoto, and Gregory Kucherov. 2014. “Using Cascading Bloom Filters to Improve the Memory Usage for de Brujin Graphs.” Algorithms for Molecular Biology 9 (1): 2. 10.1186/1748-7188-9-2.

Sun, Jingchao, Zhining Qiu, Rob Egan, Harrison Ho, Yue Li, and Zhong Wang. 2022. “Persistent Memory as an Effective Alternative to Random Access Memory in Metagenome Assembly.” BMC Bioinformatics 23 (1): 513. 10.1186/s12859-022-05052-8.

The Galaxy Community. 2024. “The Galaxy Platform for Accessible, Reproducible, and Collaborative Data Analyses: 2024 Update.” Nucleic Acids Research 52 (W1): W83–94. 10.1093/nar/gkae410.

